# 1100 Synthetic Benchmark Problems for Dynamic Modeling of Cellular Processes

**DOI:** 10.64898/2026.03.10.710893

**Authors:** Niklas Neubrand, Timo Rachel, Tim Litwin, Jens Timmer, Clemens Kreutz, Moritz Hess

## Abstract

**Motivation:** Systems biology strives to unravel the complex dynamics of cellular processes, often with the help of ordinary differential equations (ODEs). However, the sparsity of measured data and the strong non-linearity of common ODEs introduce severe numerical problems in typical modeling tasks. This gave rise to the development of many computational algorithms that must be systematically evaluated to ensure optimal method choices. Currently, the amount of well curated models for such benchmarking efforts is insufficient, as building and calibrating biologically reasonable models based on experiments requires years of work.

**Results:** We present a large-scale collection of 1100 synthetic modeling problems, generated based on the ODE systems and experimental designs of 22 published modeling problems. This is achieved by extending a recent method for simulation of time-course data for randomly generated observation functions to also include realistic measurement patterns across multiple experimental conditions. By analyzing data and model characteristics, optimization performance and parameter identifiability, we show that the synthetic problems provide both a realistic and diverse extension of the existing problem space. Hence, the synthetic collection provides a valuable resource for benchmarking in dynamic modeling.

**Availability and Implementation:** Benchmark problems and algorithm are publicly available at https://github.com/niklasneubrand/1100SyntheticBenchmarksODE and https://zenodo.org/records/14008247.

## 1 Introduction

Dynamic modeling based on ordinary differential equations (ODEs) is a fundamental technique in systems biology [1, 2]. Such mechanistic models describe biochemical reaction networks underlying cellular processes such as signaling pathways. They include unknown parameters (e.g., reaction rate constants, Hill coefficients, or initial values) that must be inferred from experimental data [3, 4]. To dissect the roles of individual system components, models are analyzed across multiple experimental conditions – for instance, with different stimuli or genetic perturbations. In each condition, distinct observables (i.e., measurable quantities such as protein concentrations or phosphorylation states) can be recorded, depending on experimental feasibility. This variety gives rise to complex data structures and intricate mappings between the observations and the underlying ODE equations [3, 4].

To manage this complexity of real-world modeling problems, several dedicated computational tool-boxes have been developed. Examples include Data2Dynamics [5], dMod [6], pyPESTO [7], and PEtab.jl [8]. These environments typically rely on standardized, human-readable specifications such as the Data2Dynamics or PEtab formats [9]. Together, they provide integrated frameworks for model simulation, parameter calibration, and uncertainty quantification, supporting both frequentist [10, 11] and Bayesian approaches [7]. These frameworks also enable higher-level modeling tasks such as model-reduction [12] or automated model selection [13].

However, despite this extensive tool support, modeling projects are often complicated by numerical challenges. Solving stiff and high-dimensional ODE systems can be computationally demanding or even unfeasible [4, 14]. Moreover, the non-linearity of ODE solutions gives rise to objective functions with non-convex regions and multiple local optima; while sparse or noisy experimental data can induce flat directions in the parameter space, termed as non-identifiability [15– As a result, parameter estimation frequently suffers from convergence issues, identifying the global optimum reliably remains challenging [3, 14].

To address these issues, it is essential to select methods and configurations that are well suited to the specific modeling problem. In practice, however, these choices are often based on experience or trial-and-error. To derive more general recommendations, systematic benchmarking across a wide range of modeling scenarios is required. While several benchmarking studies have examined particular aspects such as parameter estimation [14, 18, 19] or sensitivity computation [20], these typically rely on small sets of ODE models or simplified synthetic problems and are therefore limited in scope. To improve and standardize such efforts, a curated collection of 20 peer-reviewed models calibrated on experimental data has been published as a general benchmark resource [19]. For comprehensive benchmarking, such problems must include not only the ODE models themselves but also curated experimental datasets, observable definitions, and complete mappings between measurements and model variables. However, because both experimental research and mathematical model development are labor-intensive processes, the number of such fully specified modeling problems remains limited.

In this work, we apply synthetic data generation to scale up the existing benchmark collection. We build on a recent method for generating time-course data sets for randomly generated observables of a given ODE system [21]. To better mimic the complexities of real-world modeling problems, we extend this algorithm to work with multiple experimental conditions and develop a method to generate realistic observable patterns across experimental conditions. Central to our approach is the use of a *simulation template*, which consists of the ODE system, experimental conditions, and observation functions of an existing calibrated model. This template provides the structural and biological context from which diverse synthetic modeling problems are generated.

Using 22 published modeling problems from systems biology – including intracellular reaction networks such as JAK-STAT signaling [22–25] or population-level cell dynamics [26] – we generate a collection of 1100 synthetic modeling problems. We observe both realism and diversity in key problem characteristics. The synthetic problems reproduce the statistical distributions of the template collection while exhibiting substantial variability within each template subgroup. Furthermore, parameter optimization and identifiability analyses show that the synthetic collection extends beyond the template problems by also including more challenging cases with harder optimization landscapes and reduced parameter identifiability. With these properties, the synthetic collection provides a valuable resource for method development and systematic benchmarking of computational tools for dynamic modeling in systems biology.

## 2 Methods and Algorithm

### 2.1 Dynamic Modeling with ODEs

Throughout this work we consider biological systems, e.g. metabolic networks or signaling pathways, that can be expressed by a system of ordinary differential equations (ODEs)

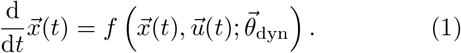

In the systems biology context, the *dynamic variables* 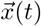 typically represent concentrations of molecular species; 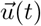 are input functions and 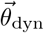the dynamic parameters (e.g. reaction rate constants or Hill coefficients). An approximate solution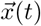can be obtained by numerical integration of the ODEs. Initial values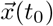are either known, derived from steady state assumptions, or estimated from the data.

To simulate different biological scenarios, e.g. genetic variations or external stimuli, parameters, initial values or input functions of the original ODEs can be replaced or modified. Each modification defines a distinct *simulation condition* and requires an independent computation of the ODE solution.

Furthermore, the biological entities described by the ODE system often cannot be measured directly. To account for this, we introduce *observation functions g*, which map internal model states to measurable quantities. Observation functions may combine dynamic variables by sums or ratios, include transformations (e.g., logarithmic), and often introduce additional *observation* parameters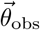 such as scaling factors, offsets, or detection limits.

Using these observation functions, we can model experimental observations *y*_*i*_ (with counting index *i* = 1, …, *N* ) by the statistical model

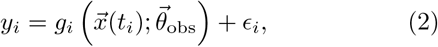

where *t*_*i*_ denotes the measurement time, *g*_*i*_ the corresponding observation function, and 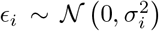the measurement noise. For simplicity, we assume independent, normally distributed residuals. Log-normally distributed residuals can be described by logarithmic transformations of both *y*_*i*_ and *g*_*i*_.

The measurement uncertainty *σ*_*i*_ is allowed to vary across data points and is often specified by an *error model* with parameters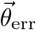, which typically combine absolute and relative error components. Importantly, the observable and error parameters are often condition-specific to account for different experimental setups, measurement techniques and batch effects.

Together, these modeling assumptions define the likelihood function 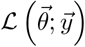which allows joint estimation Neubrand et al. 1100 Synthetic Benchmark Problems for Dynamic Modeling of Cellular Processes of all unknown parameters 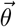by the maximum-likelihood approach

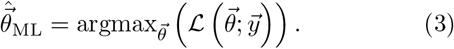

### 2.2 Formalizing Typical Data and Observable Structures

In published modeling problems observation functions and simulation conditions are carefully chosen to address specific biological questions. To describe different experimental designs, we distinguish between two types of *experimental conditions* :

- *time-course* experiment probes the dynamic behavior of the system under a single simulation condition, with measurements taken at multiple time points.
- *dose-response* experiment evaluates the system’s response to varying input levels. It consists of multiple simulation conditions (e.g., different doses) but collects measurements at a single time point.

Due to technical reasons, measurements of the same underlying system components may be described by multiple different observation functions. Common examples are condition-specific parameters or transformations. To have a common description for these related measurements, we use the term *observable* for the set of all observation functions sharing an unique subset of dynamic variables. For example, the functions *g*_1_(*x*) = *x*_1_ and *g*_2_(*x*) = *a · x*_1_ belong to the same observable, while *g*_3_(*x*) = *x*_1_ + *x*_2_ belongs to a different one, as it includes an additional dynamic variable.

In published modeling problems, we often observe structured patterns in how observables are distributed across experiments. We formalize this structure with the boolean *Experiment–Observable Matrix* (EOM), in which rows represent experiments and columns represent observables. The entry EOM_*jk*_ indicates whether a given observable *k* is measured in a particular experiment *j*. Examples for both an experimental and a synthetic modeling problem are shown in Figs. 1 A) and D), respectively.

**Figure 1.**
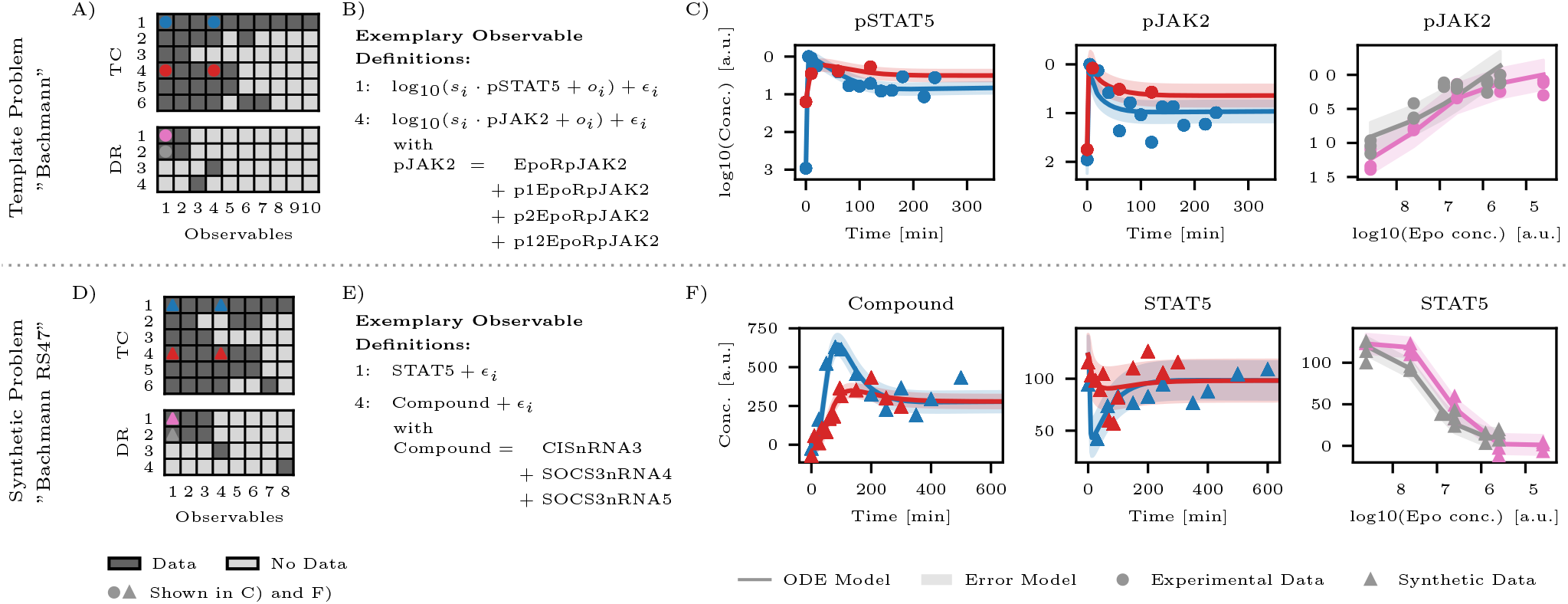
Comparison between a template problem (A-C) and a synthetic counterpart (D-F). Panels A and D show the experiment-observable matrix (EOM). The matrices are split in time-course (TC) and dose-response (DR) experiments. As shown by two examples each, the observable expressions for template (B) and synthetic (E) problems differ structurally. *s*_*i*_, *o*_*i*_, and *ϵ*_*i*_ represent experiment-dependent scaling factors, offsets, and residual errors, respectively; while all other symbols denote dynamic ODE variables *x*_*i*_(*t*) (see Eq. 1). Panels C and F show experimental (C) and synthetic (F) data for a selection of TC and DR experiments (as indicated in the EOMs) together with fitted (C) and ground-truth (F) ODE and error models. Both problems exhibit similar EOM structures, observable definitions, and data characteristics.

### 2.3 Generating Realistic Synthetic Benchmark Problems

We developed a simulation pipeline that generates realistic synthetic modeling problems based on existing modeling problems, referred to as *template problems*. Each synthetic problem is constructed in several steps:

- **Selecting Simulation Templates**: Manually choose a real-world modeling problem with known experimental structure.
- **Perturbing Dynamic Parameters**: Perturb the dynamic parameters to generate novel system behavior.
- **Generating Realistic Observable Structures**: Sample biologically plausible observables from the system states and assign them realistically to experiments.
- **Generating Realistic Synthetic Data**: Generate measurement time grids, simulate model output and add biologically calibrated noise models.

Each step introduces controlled variability while preserving biological plausibility which enables the generation of realistic and diverse benchmark problems.

#### 2.3.1 Selecting Simulation Templates

In our simulation study we use a set of 22 real-world modeling problems from systems biology as simulation templates. The models describe intracellular reaction networks, e. g. the JAK-STAT signaling pathway [22–25], or cell population dynamics [26]. All models are publicly available as examples in the Data2Dynamics toolbox. 19 problems have already been part of the experimental benchmark collection [19], and three additional suitable problems were included [27–29]. In this article, we will refer to the templates by the last name of the first author. An overview table with problem characteristics and all references is given in the supplementary material.

#### 2.3.2 Perturbing Dynamic Parameters

To generate diverse system behaviors, we perturb the dynamic parameters of the simulation template. Specifically, each calibrated parameter 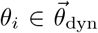 is multiplied by a random factor 2^*η*^, with *η∼* 𝒰 (*−*1, 1). This transformation is symmetrical on the logarithmic scale and introduces realistic variability in the model dynamics.

#### 2.3.3 Generating Realistic Observable Structures

To generate realistic observable structures for our synthetic benchmark problems, we build on the approach of Egert and Kreutz [21]. Their method generates realistic synthetic observation functions using a sequence of random decisions, calibrated to match the observable characteristics of 19 real-world modeling problems. The resulting observable expressions are either functions of a single dynamic variable *x*_*i*_ or sums over multiple variablesΣ_*i*_ *x*_*i*_ – termed *compound measurements*. Additionally, the expressions may include scaling, offset parameters, or log_10_ transformations. The number of observables is also randomly selected based on the ODE system dimension, with a typical ratio between the numbers of observable and dynamical variables of about 50% [21].

Neubrand et al. 1100 Synthetic Benchmark Problems for Dynamic Modeling of Cellular Processes 4 We extend this approach with realistic assignments of observation functions across multiple experimental conditions. Specifically, we generate a synthetic EOM by sampling for each synthetic observable a column from the template’s EOM (uniformly with replacement). This approach transfers the template’s measurement patterns to the synthetic problem.

To avoid generating non-informative synthetic data, observation functions are only sampled from dynamic variables with time variation in at least on experiment. After sampling the synthetic EOM, we also replace assignment of time-constant observables within each experiment to increase the time-variation in the experimental design.

We emphasize that a single observable may be realized by different observation functions in different experiments. Specifically, we define individual scale, offset, and error parameters for each experiment, even when the underlying observable is shared.

#### 2.3.4 Generating Realistic Synthetic Data

To generate data for time-course experiments, we first determine the sampling time points. For this, we use the method of Egert and Kreutz [21], which applies the Retarded Transient Function (RTF) approach [30] to analyze the dynamics of each observable. The RTF is a parametric function tailored to approximate typical ODE dynamics by combining transient and sustained dynamics. A pre-calibrated linear model then maps the RTF parameters to a realistic time grid, including number of points, time range, and spacing.

For dose-response experiments, we retain the original dose levels from the template problem but introduce variability by randomizing the sampling time. Specifically, the original measurement time is scaled by a random factor 2^*η*^, with *η ∼* 𝒰 (*−*1, 1)

Synthetic observations are generated by evaluating the observation functions at the chosen time points and adding Gaussian measurement noise:

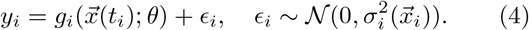

If the observable is defined on a logarithmic scale, this results in log-normal noise in the data space. The noise level 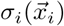is randomly drawn according to the model in [21], which was calibrated on error characteristics of 19 real-world modeling problems.

Note that scaling and offset parameters only have a minor influence on the generated data, because their ground-truth values are set to neutral values (scaling factor = 1, offset ≈1% of mean value). Therefore, these parameters only have are mostly relevant for parameter estimation tasks.

### 2.4 Evaluation Criteria

In the following, we describe the criteria and computational procedures used to evaluate the realism, complexity, and information content of the synthetic benchmark problems. We begin with basic data and model characteristics, then outline two central analysis tasks commonly encountered in dynamical modeling: multi-start parameter estimation and local identifiability analysis. Finally, we perform a principal component analysis to combine multiple metrics and to assess the structure of the full benchmark collection.

#### 2.4.1 Data and Model Characteristics

To characterize each benchmark problem, we define a set of key characteristics describing both the data and the underlying statistical model. As high-level indicators of problem size and complexity, we report the number of observables, estimated parameters, and data points.

We quantify the overall noise level of each modeling problem by the median relative error. The relative error for each data point is defined as

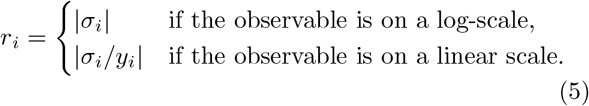

For synthetic problems, *y*_*i*_ and *σ*_*i*_ refer to the generated data values and ground-truth error parameters. For template problems, *y*_*i*_ are experimental measurements, and *σ*_*i*_ are either measured uncertainties or fitted error models.

In addition, we report the median measurement time to reflect both the time-scales of the system and the data sampling.

#### 2.4.2 Multi-Start Parameter Estimation

To explore the likelihood landscape and investigate the performance of parameter estimation, we conduct a global optimization analysis using multi-start local optimization. For each benchmark problem, we uniformly sample 100 random initial parameter vectors within the bound constraints. For the template the parameter scale (linear or logarithmic) and bounds are predetermined. For synthetic problems, we use the template bounds and scales for dynamic parameter. For the novel generated error, scale and offset parameters we choose a logarithmic scale with bounds extending two orders of magnitude in both directions.

For each initial parameter guess, we independently perform local optimization of the likelihood function using the trust-region reflective algorithm [31]. The ODE solver uses absolute and relative tolerances of 10^*−*8^ to ensure numerical accuracy, while the maximum number of solver steps is set to 10^5^.

Neubrand et al. 1100 Synthetic Benchmark Problems for Dynamic Modeling of Cellular Processes 5 Optimization runs may terminate in one of three ways: by ODE solver failure, by fulfilling the optimizer’s local convergence criterion, or by reaching the optimizer’s iteration limits without local convergence. As performance metrics of the optimization we consider the numbers of failed and of locally converged optimization runs.

Additionally, we report the frequency at which the global optimum has been found by the multistart approach. This can be seen as a measure for the complexity of the optimization problem. We define it by counting the number of optimization runs with final objective function value *−*2 log(ℒ ) within a tolerance of 10^*−*2^ from the best observed value.

#### 2.4.3 Local Identifiability Analysis

To assess the information content of the data sets, we examine which parameters are structurally or practically non-identifiable. For this purpose, we apply the Identifiability Test by Radial Penalization (ITRP) [32], a method that detects flat directions in the likelihood function through penalized local optimization. When such a direction is found, the most strongly aligned parameter is classified as non-identifiable and excluded from further searches. The method can be applied repeatedly until all non-identifiable parameters are found.

The ITRP algorithm requires initialization at a local optimum. Therefore, we performed local optimization as described above for all modeling problems. For synthetic problems, we use the ground truth parameters as an initial parameter vector. Although the template problems are already optimized, we run the optimization algorithm to test the ODE solver and optimizer configuration.

The test is then applied with its default configuration: a penalty radius of 1, five radial directions tested per iteration, and a flatness threshold of 10^*−*3^. The maximum number of ODE solver steps was set to 10^6^. All other settings are consistent with the multi-start parameter estimation described above.

The statistical models have different kinds of parameters: the dynamic parameters (e.g. rate constants and initial values) determine the dynamics of the ODE system, while all remaining parameters are responsible for the mapping from the ODE system to the data by observable and error models. For our problem-generating algorithm this distinction is especially important, since the ODE system and all dynamic parameters are taken directly from the template problem, while the observable and error parameters are introduced by generation and condition-assignment of the observables. Thus, we define the identifiability ratio for three parameter sets: “all parameters”, the “dynamic parameters” and the remaining “non-dynamic parameters”.

#### 2.4.4 Principal Component Analysis

To visualize the structure of the benchmark collection, we perform a principal component analysis (PCA) on 14 numerical features that capture key aspects of problem size, experimental design, parameter identifiability, and optimization behavior. The selected features are:

- **Template Characteristics:** numbers of dynamic variables, estimated dynamic parameters, time-course experiments, and dose-response experiments.
- **Problem Characteristics (see Sec. 2.4.1):** numbers of observables, estimated parameters, and data points; median relative error and measurement time.
- **Parameter Estimation (see Sec. 2.4.2):** fractions of failed and converged runs, and frequency of global optimum recovery.
- **Parameter Identifiability (see Sec. 2.4.3):** identifiability ratios for dynamic and non-dynamic parameters.

Note that the first four features are defined by the template problem alone and are therefore shared across all corresponding synthetic problems. The remaining features vary at the level of individual benchmark problems. To ensure numerical stability and comparability, all count-based features *x* are log-transformed using log_10_(*x*+10^*−*6^) and then standardized. Ratio-based metrics are standardized directly without transformation. The PCA is conducted on the full benchmark set, including both template and synthetic problems. Benchmark problems with missing values – e.g. due to failed identifiability analysis – need to be excluded. Importantly, exclusion of a template problem does not necessarily imply exclusion of derived synthetic problems.

## 3 Results

### 3.1 Exemplary Comparison: Template vs. Synthetic

To illustrate the similarity between real and synthetic problems, we compare the template problem “Bachmann” [22] with one of its derived synthetic problems, “Bachmann RS47”. The template problem comprises 541 data points for 10 observables and 115 estimated parameters, while the synthetic problem includes 492 data points for 8 observables and 69 parameters.

Figure 1 provides a side-by-side comparison of the two problems. The experiment-observables matrices in the left columns (panels A and D) display similar measurement patterns across experiments. This showcases that most columns of the synthetic EOM are sampled from the template EOM without modification.

We emphasize, that the columns in the template and synthetic EOM do not correspond to the same observable expressions. In the middle column (panels B and E) we present two exemplary observable definitions for each problem. In contrast to the template, the synthetic observables do not include any transformations or unknown parameters. Still, both problems contain both individually observed dynamic variables, as well as compound measurements of similar complexity. Thus, the synthetic observable complexity seems comparable to real-world models.

The last column (panels C and F) show selected time-course and dose-response data sets for these observables. The plots cannot be compared directly, due to different observable definitions and randomly perturbed dynamic parameters of the synthetic problem. Nonetheless, the synthetic data is similar to the template data in key qualitative features: the number of data points per experiment, the sampling density, and the relative noise levels.

### 3.2 Data and Model Characteristics

To assess the realism and diversity of the generated benchmark problems, we computed five key characteristics: number of observables, number of parameters, number of data points, median relative error, and median measurement time.

Looking at the collection as a whole, the median synthetic problem contains 9 dynamic variables, 4 observables, 106 data points, and 37 estimated parameters – along with a median relative error of 13% and a median measurement time of 74 s (see Fig. 2). These values fall well within the range of realistic calibration scenarios. The pooled synthetic distributions, visualized by stylized boxplots in the panel backgrounds, are symmetrical, contain few outliers, and span ranges similar to those of the template problems. This supports the realism and plausibility of the synthetic modeling problems.

**Figure 2.**
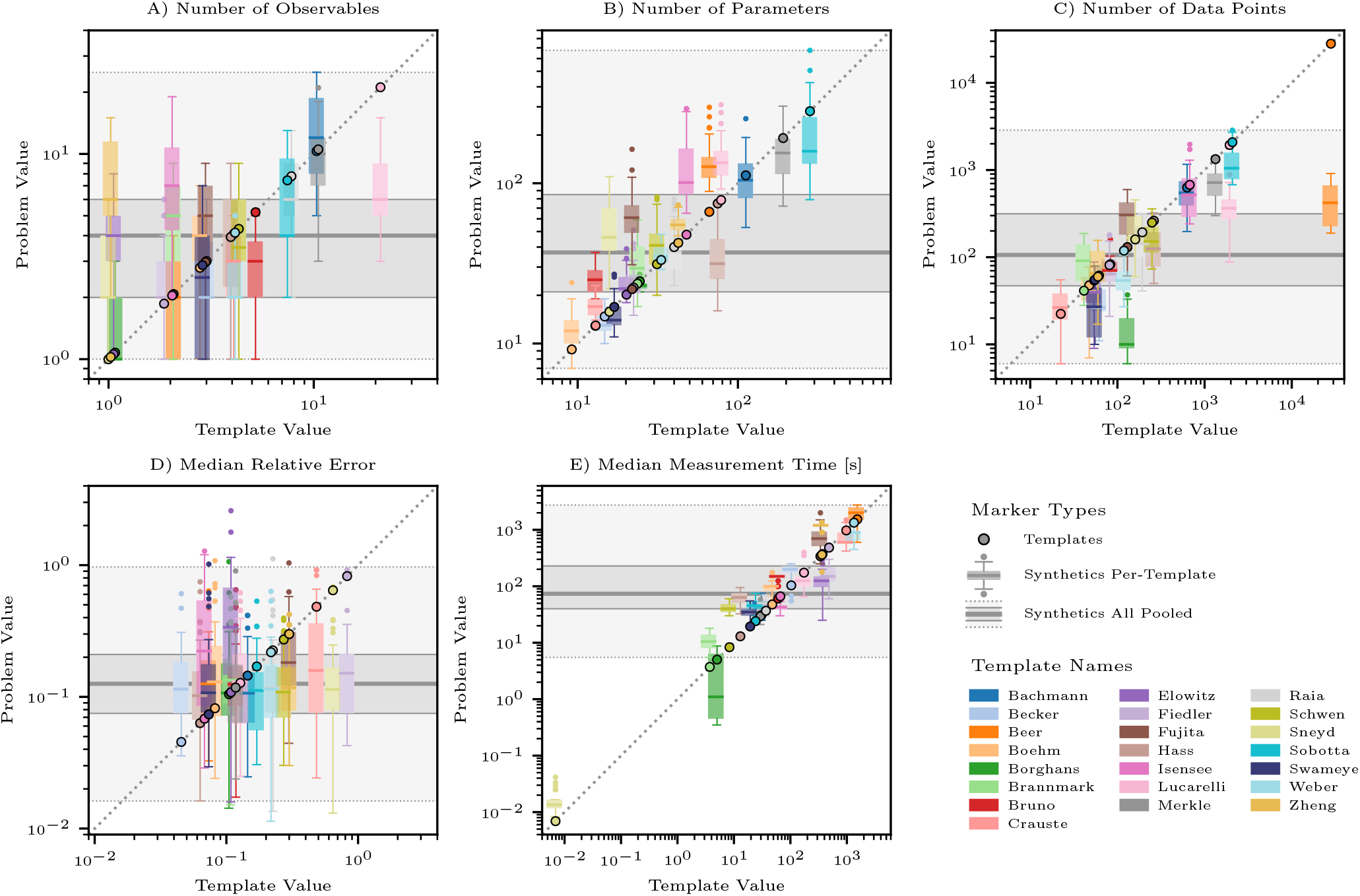
Comparison of synthetic and template problem characteristics. Each panel shows the values for the template problem (horizontal axis) versus the distributions across its 50 synthetic counterparts (vertical axis), summarized as colored boxplots. Small markers show outliers of the per-template distributions. Colors indicate the template name (e.g., Bachmann [22], Becker [33]). Slight horizontal jitter improves readability. The background shows pooled distributions across all synthetic problems as gray boxplots: solid lines for median and quartiles, dotted lines for whiskers (1.5×IQR), and shaded areas for visual guidance.

Stratified per-template distributions are generally symmetrical with few outliers and exhibit substantial internal variability. As a result, the distributions of different templates frequently overlap. This indicates that the synthetic problems do not form isolated clusters around their respective templates, but instead constitute a unified and collection that spans a broad and continuous problem space.

The collection of synthetic problems also includes combinations of model characteristics not observed in the original template set. This indicates that our algorithm substantially enhances the problem diversity.

### 3.3 Multi-Start Parameter Estimation

To illustrate a typical modeling task, we performed multi-start parameter estimation on all synthetic and template problems. For each problem, 100 optimization runs were conducted with randomly initialized parameter values. Figure 3 shows the distributions of three key metrics: the number of failed runs, locally converged runs, and global optimum hits. In addition to boxplots, individual values are shown as strip plots to highlight the difference in group sizes between synthetic and template problems.

**Figure 3.**
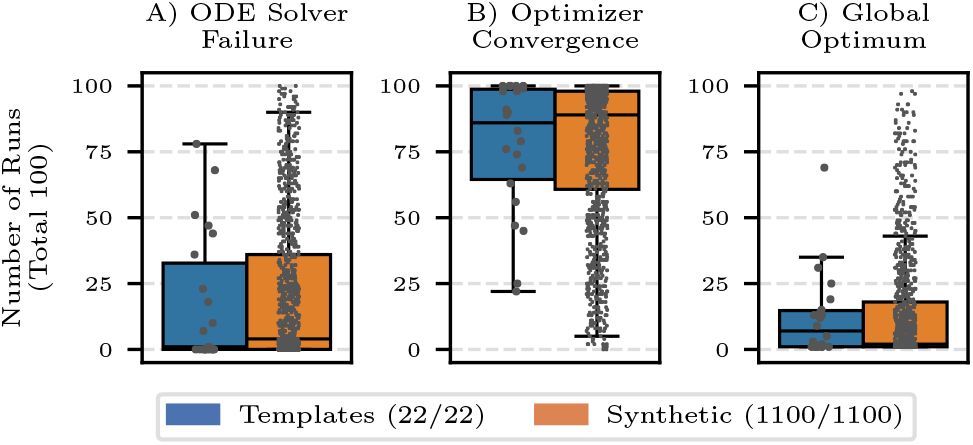
Counting statistics for multi-start parameter estimation. A) Number of optimization runs with ODE solver failure. B) Number of runs reaching local convergence criterion. C) Number of runs that recovering the best objective value. In each panel, blue boxplots show results for template problems, and orange boxplots for synthetic problems. Individual values are overlaid as strip plots. The legend indicates that all problems were included in the analysis. Synthetic benchmarks exhibit substantial optimization diversity, while the majority can be optimized sufficiently well.

Most synthetic problems are reasonably well optimizable. Specifically, 75% of problems had fewer than 36 failures and more than 60 converged runs out of 100. These thresholds correspond to the boxplot quartiles shown in panels A and B. For the template problems, we observed similar performance, with 75% of problems below 33 failures and above 64 converged runs. Notably, the synthetic distribution has a longer tail toward “hard-to-optimize” cases, where the majority of runs fail due to ODE solver errors. This reflects that the generation process not only replicates typical optimization challenges but also produces more varied – and sometimes less polished – modeling problems.

To assess the complexity of the optimization landscape, we also counted how often the global optimum was recovered – defined as reaching the best observed objective value within a tolerance of 10^*−*2^. For 43% of synthetic and 31% of template problems, the best value was found only once, suggesting potentially complex optimization landscapes or numerical instabilities. Due to the large number of synthetic problems, the distribution again shows a long tail, this time toward “easier” problems with high recovery rates of the global optimum. However, the upper quartile remains low (18/100 for synthetic and 15/100 for template problems), indicating that most problems exhibit non-trivial optimization landscapes.

In summary, the synthetic problems reflect a realistic spectrum of optimization behavior, including both well-behaved and challenging cases. This confirms that the benchmark set achieves its goal of combining realism with diversity in optimization difficulty.

### 3.4 Local Identifiability Analysis

To evaluate the information content of the benchmark problems, we analyzed parameter identifiability using the Identifiability Test by Radial Penalization (ITRP) [32]. This method searches for flat directions in the likelihood landscape to identify structurally or practically non-identifiable parameters.

The test could be successfully performed for 21 of the 22 template problems and 993 of the 1100 synthetic problems. Most failures stemmed from ODE solver issues during local optimization, likely due to unadjusted solver settings that were kept fixed for comparability.

Figure 4 shows the distribution of identifiable parameter ratios. When considering all parameters (panel A), synthetic problems show reduced identifiability compared to templates, with a median of 87% versus 93%.

**Figure 4.**
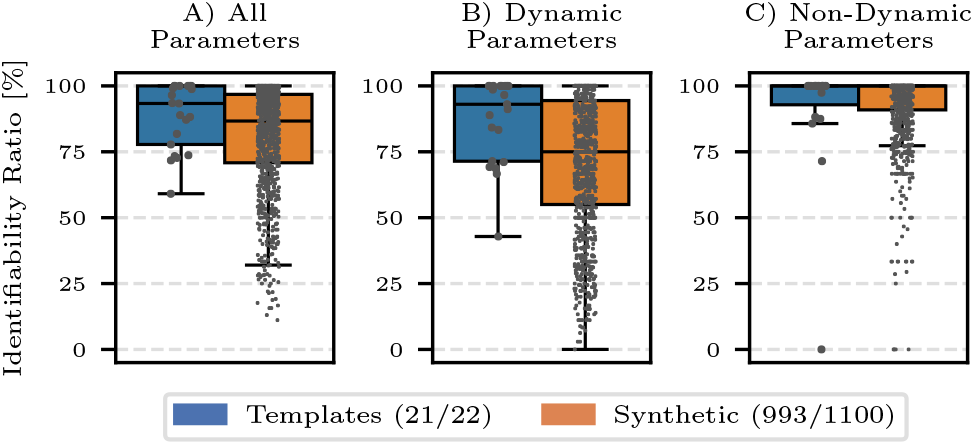
Ratio of identifiable parameters for different parameter subsets. In each panel, blue boxplots show results for template problems, and orange boxplots for synthetic problems. Individual values are overlaid as strip plots. The legend indicates the numbers of problems with successful analysis results. Synthetic problems generally generate more non-identifiable parameters, especially for the dynamic parameters.

Separation in parameter subsets, reveals that this difference clearly arises from the dynamic parameters (panel B). Here, synthetic problems have a wider distribution, with quartiles ranging from 55% to 94%, whereas template problems range from 71% to 100%. In contrast, non-dynamic parameters (panel C) are consistently well-identifiable in both groups, with both medians at 100% and lower quartiles at 91% for synthetic and 93% for template problems.

These results suggest that reduced identifiability in synthetic problems is primarily due to limited observability of dynamic system components. This is consistent with the randomized construction of observation functions, which may leave some states insufficiently constrained.

### 3.5 PCA Projection of Problem Space

To complement the detailed evaluation metrics, we performed a principal component analysis (PCA) to obtain a low-dimensional representation of the benchmark collection. Of the 1122 problems, 1010 (19 templates and 991 synthetics) had complete data for all 14 selected characteristics (see Sec. 2.4.4) and were included in the analysis. Note, that exclusion of a template problem does not imply exclusion of the derived synthetic problems.

Figure 5 shows a scatterplot of the first two principal components, which together explain 49.8% of the total variance. Many synthetic problems are located close to their corresponding templates, indicating successful preservation of core characteristics. At the same time, the synthetic problems exhibit greater variability, leading to substantial overlap between template-specific regions and filling in much of the space between them. As a result, the synthetic collection forms a nearly continuous distribution which only includes two larger gaps between otherwise connected clusters.

**Figure 5.**
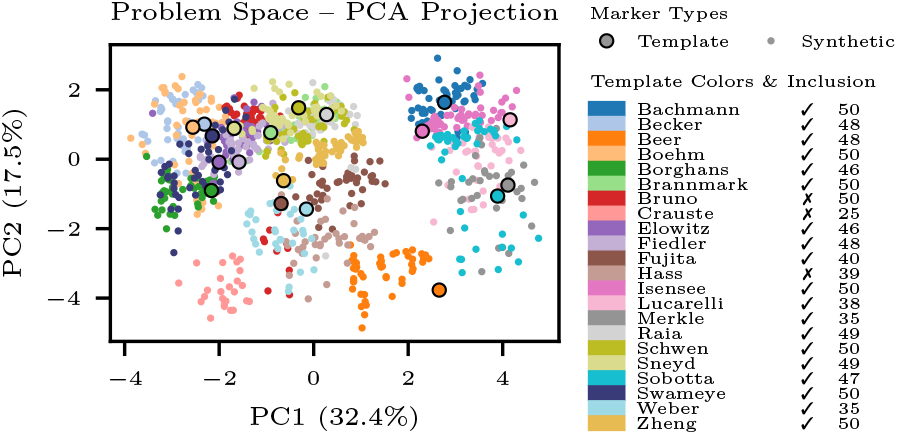
PCA projection of the problem space based on 14 problem characteristics. Template problems are shown as edge-highlighted circles and synthetic problems as small dots; colors indicate the corresponding template. Problems with incomplete feature vectors are excluded form the PCA. The legend reports for each template, its inclusion status (✓ or ✗) and the numbers of included synthetic problems.

## 4 Discussion

In this work, we introduced a large-scale collection of 1100 synthetic benchmark problems for dynamic modeling of cellular processes. Each problem is based on one of 22 template modeling problems that provide the biological context and modeling structure in form of ODE equations, definitions of experimental conditions and their relationship to measured observables. Our algorithm generates synthetic data sets for randomly sampled observables with realistic measurement patterns across experimental conditions. This results in novel synthetic modeling problems. Both the synthetic benchmark collection and the problem-generation algorithm are made publically available to the scientific community.

### 4.1 Realism and Diversity

A central strength of our approach is combining the realism of experimental models with the scalability of synthetic data generation. The two key evaluation criteria for the resulting benchmark collection are realism and diversity: Are the synthetic problems representative of real-world modeling tasks? And do they go beyond the templates to add value?

To evaluate these aspects, we analyzed the benchmark problems in terms of size, complexity, noise level, optimization performance, and parameter identifiability [32]. The realism of the synthetic problems is supported by multiple observations. The underlying ODE systems and experimental designs are based on experimentally derived biological models. The synthetic summary statistics (e.g., number of data points, parameters, noise levels) fall into realistic ranges and typically encompass the template values. Moreover, many synthetic problems exhibit similar optimization performance and parameter identifiability.

Beyond realism, the benchmark collection demonstrates substantial diversity. The synthetic distributions are continuous and wide, often exceeding the ranges of the templates. The multi-start optimization results and identifiability tests reveal that the synthetic problems cover a wide spectrum – from simple, well-constrained models to highly challenging or partially unidentifiable ones. While the latter may initially appear unrealistic, these problems can be interpreted to reflect common intermediate stages in real-world modeling workflows, where model refinement (e.g. in terms of model selection of reduction) is still ongoing.

Together, these observations show that the synthetic benchmarks not only replicate known problem types but also enrich the problem space by including new combinations of characteristics and varying degrees of complexity.

### 4.2 Applications

The benchmark collection is intended as a general-purpose resource for evaluating methods in dynamic modeling. Typical tasks in the modeling workflow involve parameter estimation, identifiability and uncertainty analysis. Thanks to the randomized experimental designs with only partial observations they can be particularly useful also for model selection and reduction. As seen by the results of the local identifiability analysis, many of the synthetic benchmarks include non-identifiable parameters. Removing these non-identifiabilities could be formulated as both model reduction and model selection tasks. Additionally, individual problems or subsets can serve as test cases in the development of new computational methods.

### 4.3 Limitations and Outlook

While the template-based simulation approach ensures high realism, it also introduces a nested structure in the synthetic benchmark collection when multiple synthetic problems are generated from the same template problem. Our analysis of general problem characteristics and the PCA projection of the problem space, showed a large variability of the generated problems. While there are clusters remaining, we mostly see interconnected and overlapping per-template distributions which supports a reduced impact of the nesting. Nevertheless, this structure is inherent to the synthetic benchmark collection and should always be considered when analyzing benchmark results.

In the future, the variability of generated problems could be further increased by extending the problem-generating algorithm. For instance the definitions of experimental conditions could be generated synthetically or the ODE equations could be randomly manipulated (e.g. by adding or removing reactions). Moreover, the problem generating algorithm could be modified to give the user control over certain problem characteristics. This way one could intentionally generate specific synthetic problems instead of relying on randomization results.

### 4.4 Conclusion

We introduced a large collection of 1100 realistic and diverse synthetic benchmark problems for dynamic modeling. By densely populating and extending the problems space defined by published models, the collection serves as a valuable resource for systematic method development, evaluation, and benchmarking in systems biology.

## 5 Data availability statement

All modeling problems (template and synthetic), the problem-generation algorithm and J. all analysis scripts are available on GitHub at F. https://github.com/niklasneubrand/1100SyntheticBenchmarks ODE and Zenodo [34] at https://zenodo.org/records/14008247. Modeling problems are shared in their native Data2Dynamics file format [3, 5]. We also provide them as PEtab [9] files. But not not all problems can correctly represent in this format. More details are given in the supplementary material.

## 6 Competing interests

No competing interest is declared.

## 7 Author contributions statement

N.N. and C.K. conceptualized the project. N.N. and T.R. worked on code implementation. N.N. performed all computations and analyses. N.N. and T.L. wrote the initial manuscript draft. All authors reviewed the final version. C.K., M.H. and J.T. supervised the project.

## 8 Acknowledgments

Funded by the Deutsche Forschungsgemeinschaft (DFG, German Research Foundation) – Project-ID 499552394 – SFB 1597.

## Notes

https://github.com/niklasneubrand/1100SyntheticBenchmarksODE

https://zenodo.org/records/18931839

### Competing Interest Statement

The authors have declared no competing interest.

