## Supplementary Material for "1100 Synthetic Benchmark Problems for Dynamic Modeling of Cellular Processes"

Niklas Neubrand[1,*], Timo Rachel[1], Tim Litwin[1,2], Jens Timmer[2,3,4], Clemens Kreutz[1], Moritz Hess[1]

**1** Institute of Medical Biometry and Statistics, Faculty of Medicine and Medical Center, University of Freiburg, Germany
**2** Center for Data Analysis, Modeling and AI (FDMAI), University of Freiburg, Germany
**3** Institute of Physics, University of Freiburg, Germany
**4** Centre for Integrative Biological Signalling Studies (CIBSS), University of Freiburg, Germany

### 1 Experimental Template Problems

#### 1.1 Template Overview

Table 1 lists all templates together with their problem characteristics and the original references.

| Template | States | Experiments TC | DR | Observables | Parameters Dyn | All | Data Points | Median Error [%] | Median Time | Ref |
|---|---|---|---|---|---|---|---|---|---|---|
| Bachmann | 25 | 6 | 4 | 10 | 29 | 115 | 541 | 15 | 30 s | [1] |
| Becker | 6 | 1 | 0 | 3 | 9 | 14 | 72 | 5 | 2 min | [2] |
| Beer | 4 | 19 | 0 | 2 | 70 | 72 | 27132 | 8 | 30 min | [3] |
| Boehm | 8 | 1 | 0 | 3 | 6 | 9 | 48 | 9 | 45 s | [4] |
| Borghans | 3 | 1 | 0 | 1 | 20 | 23 | 111 | 11 | 5 s | [5] |
| Brannmark | 9 | 2 | 4 | 2 | 14 | 22 | 43 | 12 | 5 s | [6] |
| Bruno | 7 | 6 | 0 | 5 | 13 | 13 | 77 | 11 | 1 min | [7] |
| Crauste | 5 | 1 | 0 | 4 | 12 | 12 | 21 | 44 | 13 min | [8] |
| Elowitz | 8 | 1 | 0 | 1 | 16 | 19 | 58 | 11 | 5 min | [9] |
| Fiedler | 6 | 3 | 0 | 2 | 12 | 22 | 72 | 82 | 6 min | [10] |
| Fujita | 9 | 6 | 0 | 3 | 19 | 22 | 144 | 27 | 8 min | [11] |
| Hass | 9 | 4 | 4 | 4 | 0 | 72 | 221 | 7 | 15 s | [12] |
| Isensee | 25 | 15 | 16 | 2 | 34 | 46 | 687 | 7 | 1 min | [13] |
| Lucarelli | 33 | 11 | 1 | 21 | 72 | 84 | 1755 | 12 | 3 min | [14] |
| Merkle | 23 | 16 | 11 | 11 | 43 | 199 | 1137 | 13 | 30 s | [15] |
| Raia | 14 | 4 | 0 | 8 | 18 | 39 | 205 | 25 | 35 s | [16] |
| Schwen | 11 | 7 | 0 | 4 | 13 | 30 | 286 | 26 | 8 s | [17] |
| Sneyd | 6 | 8 | 0 | 1 | 14 | 15 | 135 | 72 | 8 ms | [18] |
| Sobotta | 13 | 32 | 21 | 8 | 23 | 260 | 2222 | 18 | 20 s | [19] |
| Swameye | 9 | 1 | 0 | 3 | 10 | 17 | 46 | 7 | 15 s | [20] |
| Weber | 7 | 3 | 0 | 4 | 26 | 36 | 135 | 23 | 24 min | [21] |
| Zheng | 15 | 1 | 0 | 1 | 45 | 46 | 60 | 29 | 8 min | [22] |

**Table 1:** Overview of all experimental template problems, including key model characteristics and original references. "TC" and "DR" refer to time-course and dose-response experiments, respectively. The parameter counts include only those estimated from experimental data, with "Dyn" indicating the subset of dynamic parameters. Median relative errors are reported as percentages. "Median Time" refers to the median measurement times of the data.

#### 1.2 Implementation Details

The exact implementations of the all template problems can be found in our GitHub repository at https://github.com/niklasneubra

For most templates, the implementation is directly taken from the "Examples" folder of the Data2Dynamics toolbox [23, 24]. However, for some templates we made slight implementation changes due to technical reasons. These changes are described in the following.

##### 1.2.1 Becker

In the modeling problem of Becker et al. [2] parameters are estimated jointly for an ODE system for Erythropoetin (Epo) receptor internalization and a Michaelis-Menten model for the binding kinetics. Since our problem generating algorithm currently only supports modeling problems with a single ODE system, we removed the Michaelis-Menten model to create the simulation template.

##### 1.2.2 Merkle

The modeling problem of Merkle et al. [15] was constructed to identify cell-type specific differences in Epo-induced JAK2-STAT5 signaling. It was originally implemented in the Data2Dynamics toolbox by utilizing two very similar ODE systems – each representing one cell type. To align this with our problem generation algorithm, we modified this implementation to use a single ODE system only. The cell-type specific variations of this ODE system are represented by parameter replacements in the definitions of additional experimental conditions. To summarize, we transformed the modeling problem to another equivalent representation, without altering the parameter estimation task.

##### 1.2.3 Swameye

The ODE system of Swameye at al. [20] uses a spline function to model the input of phosphorelated Epo receptors to the cell. While the time-points of the spline knots were predetermined by the authors, the amplitudes are free parameters that need to be estimated from experimental data.

In our problem-generating algorithm all free parameters – including the spline amplitudes – are randomly perturbed. We observed that this randomization induced divergent behavior of the input function for larger simulation times. Therefore, we decided to further constrain the shape of the input spline by adding additional splines with fixed time-points and amplitudes. Specifically, we replace the original 5-knot spline by a 10-knot spline to force the input function towards small values for large simulation times. This way, we keep the original set of dynamic parameters, while allowing some controlled randomization of the input dynamics.

### 2 Code Implementation Details

#### 2.1 Working with Data2Dynamics

The simulation pipeline is implemented in MATLAB and utilizes the Data2Dynamics toolbox [23, 24].

Template problems can be provided in the Data2Dynamics file formats. Our distinction between time-course and dose-response experiments is based on the model implementation provided by the user. Data definition files (`data.def`) with identifier `PREDICTOR-DOSERESPONSE` are interpreted as a dose-response experiment.

The benchmark problem problems can easily be loaded with the Data2Dynamics toolbox by navigating to the respective problem folder and running the commands: `arInit; Setup`. Then, any analysis implemented in Data2Dynamics code can be performed.

#### 2.2 Support for PEtab

Template problems can also be provided in the interoperable PEtab format [25] and then be translated to Data2Dynamics files using the import function `arImportPEtab`.

Since the PEtab format does not include an identifier for dose-response experiments, all simulation conditions are by default interpreted as time-course experiments. This can be changed by manual modifications in the imported Data2Dynamics files.

We also provide the benchmarks in PEtab files for usage outside of the Data2Dynamics toolbox. Unfortunately, the files exported from the Data2Dynamics toolbox use non-standard mathematical expressions in the condition definitions. This problem affects 6 of the 22 template problems and the corresponding derived synthetic benchmarks.

With the upcoming release of PEtab v2, these kinds of condition expressions are expected to be officially supported and this limitation could be removed.

#### 2.3 Computational Robustness

The template problems are diverse in computational complexity and modeling features – such as pre-equilibration, input functions, or error parameter estimation. This imposes high requirements for versatility and robustness of our problem-generation algorithm. During its development we improved it iteratively to catch a large portion of typical errors.

With the current version of the algorithm, all 50 problem generation runs terminated successfully for 18 out of 22 template problems. For the remaining 4, occasional numerical issues when solving the underlying ODE systems required restarting the problem-generation algorithm with different random seeds. The most problematic case was the template based on the model by Isensee et al. [13], which required 67 runs to generate 50 valid synthetic problems (success rate of 75%).

In total, 1,136 problem-generation runs were performed to obtain 50 successful runs for each of the 22 templates, resulting in an overall success rate of 97%.

[19] Svantje Sobotta, Andreas Raue, Xiaoyun Huang, Joep Vanlier, Anja Jünger, Sebastian Bohl, Ute Albrecht, Maximilian J. Hahnel, Stephanie Wolf, Nikola S. Mueller, Lorenza A. D'Alessandro, Stephanie Mueller-Bohl, Martin E. Boehm, Philippe Lucarelli, Sandra Bonefas, Georg Damm, Daniel Seehofer, Wolf D. Lehmann, Stefan Rose-John, Frank van der Hoeven, Norbert Gretz, Fabian J. Theis, Christian Ehlting, Johannes G. Bode, Jens Timmer, Marcel Schilling, and Ursula Klingmüller. Model Based Targeting of IL-6-Induced Inflammatory Responses in Cultured Primary Hepatocytes to Improve Application of the JAK Inhibitor Ruxolitinib. *Frontiers in Physiology*, 8, October 2017.

[25] Leonard Schmiester, Yannik Schälte, Frank T. Bergmann, Tacio Camba, Erika Dudkin, Janine Egert, Fabian Fröhlich, Lara Fuhrmann, Adrian L. Hauber, Svenja Kemmer, Polina Lakrisenko, Carolin Loos, Simon Merkt, Wolfgang Müller, Dilan Pathirana, Elba Raimúndez, Lukas Refisch, Marcus Rosenblatt, Paul L. Stapor, Philipp Städter, Dantong Wang, Franz-Georg Wieland, Julio R. Banga, Jens Timmer, Alejandro F. Villaverde, Sven Sahle, Clemens Kreutz, Jan Hasenauer, and Daniel Weindl. PEtab—Interoperable specification of parameter estimation problems in systems biology. *PLOS Computational Biology*, 17(1):e1008646, January 2021.
